# Event Extraction from Biomedical Literature

**DOI:** 10.1101/034397

**Authors:** Abdur Rahman M.A. Basher, Alexander S. Purdy, Inanç Birol

## Abstract

The breadth and scope of the biomedical literature hinders a timely and thorough comprehension of its content. PubMed, the leading repository for biomedical literature, currently holds over 26 million records, and is growing at a rate of over 1.2 million records per year, with about 300 records added daily that mention ‘cancer’ in the title or abstract. Natural language processing (NLP) can assist in accessing and interpreting this massive volume of literature, including its quality NLP approaches to the automatic extraction of biomedical entities and relationships may assist the development of explanatory models that can comprehensively scan and summarize biomedical articles for end users. Users can also formulate structured queries against these entities, and their interactions, to mine the latest developments in related areas of interest. In this article, we explore the latest advances in automated event extraction methods in the biomedical domain, focusing primarily on tools participated in the Biomedical NLP (BioNLP) Shared Task (ST) competitions. We review the leading BioNLP methods, summarize their results, and their innovative contributions in this field.

## Introduction and Background

As the number of publications in the biomedical domain grows, the cost of retrieving relevant information and inferring scientific knowledge from this expanding corpus becomes a challenging task [1]. This problem has triggered interdisciplinary research at the interface between the biomedical and natural language processing domains (BioNLP) to automate the collection, formatting and curation of biological information. Though not mature enough yet, BioNLP research has already yielded some specialized tools that can go through documents to label biomedical entities, such as genes and proteins, with measurable accuracy. Supplementing to the recent surveys [2, 3, 4, 5, 1, 6], in this paper we review proposed solutions to the event extraction (EE) problem developed within the context of community-wide efforts, focusing primarily on the biennial BioNLP Shared Task (ST) competitions. We offer an independent survey of leading methods, and categorize them based on how they are conceptualized and implemented.

***Event extraction*** refers to the task of recovering structured representations of biological events from text with associated attributes and properties, which are often characterized by complex, nested argument structures involving several entities or recursively embedded relations. It is one of the unit operations in programmatic identification of associations among entities and other information in textual documents. The utility of EE include many applications in the biomedical domain [7, 8, 1]. For example, many translational genomics initiatives seek to incorporate genome-based tests into standard clinical care [9]. These tests often look for specific biomarkers that would allow physicians to develop personalized treatment plans for their patients. However, a list of mutations is not enough on its own; the results need to be correctly interpreted in the proper context, as presented in the medical literature. Event extraction systems provide the opportunity to extract accurate context of the observed mutations to cancer and treatment, as well as the opportunity to generate new hypotheses by discovering and assessing novel relationships among entities in the literature and genomic data.

The initial step in event extraction is ***named entity recognition*** (NER). NER involves delimiting the textual boundaries that constitute an entity’s name, and classifying the entity according to its associated, predefined classes [6]. Biomedical entities include gene and protein names, biological structures, diseases, and drug names. These entities are typically processed into their unique and unambiguous representations through a process called ***normalization***. In text, a particular concept can be referred in various ways, which are called ***term variants***. For example, “TIF2”, “TIF-2”, “transcription intermediary factor-2” and “transcriptional intermediate factor 2” all denote the same entity. The normalization process automates the grouping of term variants into an equivalence class, which is derived from external knowledge bases. In BioNLP-ST competitions, the entities are already provided and the normalization step is employed to increase the performance of event extraction.

Following named entity recognition task, relationships that capture the logical connections among the detected entities are determined. Automating the process of identifying relationships is an important prerequisite to the computational understanding of biological processes. Earlier biomedical event extraction efforts focused on extracting simple, pairwise relations between biomedical entities, such as protein-protein interactions (PPI) [10], gene-disease relationships [11, 12], and drug-drug interactions (DDI) [13, 14]. However, these binary relations alone are not sufficient to deeply capture biomedical phenomena, indicating the need for better representations to improve extraction of complex relations or events [15].

To address the aforementioned problem, [16] suggested the GENIA event annotation schema to represent dynamic biological processes, which often involve a change in the properties, locations or interactions between biomedical entities, such as proteins, cells and chemicals. Additionally, the schema defines attributes associated with events including polarity, negation and speculation that reshape the contextual information anchored with an event. These attributes are vitally important [17], as for example negation reverses the meaning of an event, and negation of a candidate argument prevents the construction of an event with this argument.

The problem of event extraction can be roughly decomposed into two sub-tasks - which could be learned jointly or sequentially (see “Event Extraction Methodologies” section): ***trigger detection*** and ***argument detection***. The first subtask, trigger detection, involves identifying explicit content-specific textual terms in a sentence that indicate the occurrence of an event. These textual units, or ***trigger words***, usually take the form of verbs (e.g., regulates, phosphorylates) or nominalized verbs (e.g., regulation, phosphorylation) [10]. Trigger detection is a domain-specific task, where the semantics and number of event-types may vary greatly between domains. For example, in the BioNLP 2013 Genia (GE) ST there were 13 event types (gene expression, protein catabolism or modification, regulation, etc. See Table 1 in [18]). The second subtask is that of argument detection, which involves identifying the participants of an event along with their associated semantic roles to fully construct an event. These arguments typically consist of other entities (e.g., proteins) or events that comprise nested structures. Additionally, it is not surprising that the same entity might be formed in different events due to the nature of events. The associated semantic categories of these arguments are drawn from a fixed ontology of semantic types (e.g., theme and cause roles). These two subtasks may be followed by postprocessing steps to transform the detected triggers and arguments into valid structures, as defined by some task-specific guidelines.

**Table 1.** Summary of event extraction corpora

| Corpus | Number of documents | Domain(s) | Annotation Schema | Reference Ontologies & DBs | Number of annotations |
| --- | --- | --- | --- | --- | --- |
| Genia Event (GE) 2011 [30] | 1210 abstracts, 14 full-text | humans bloods transcription factors | 2 entity types, 9 event types | GENIA Event ontology | 21616 protein, 24967 events |
| Genia Event (GE) 2013 [18] | 34 full-text | humans bloods transcription factors | 2 entity types, 13 event types | GENIA Event ontology | 12068 protein, 762 entities, 9364 events, 535 coreference |
| Epigenetics and Post-translational Modifications (EPI) [31] | 1200 abstracts | epigenetics and post-translational modifications | 2 entity types, 14 event types | GENIA Event ontology | 15190 protein, 3714 events, 369 modifications |
| Infectious Diseases (ID) [32] | 30 full-text | two-component regulatory systems | 5 entity types, 10 event types | GENIA Event ontology | 12740 entity, 4150 events, 214 modifications |
| Cancer Genetics (CG) [33] | 600 abstracts | cancer biology | 18 entity types, 40 event types | CARO, taxonomy DBs, CL, GO, gene/protein DBs, ChEBI | 21683 entities, 917 relations, 17248 events, 1326 modifications |
| Pathway Curation (PC) [34] | 525 abstracts | highly annotated pathway models | 4 entity types, 23 event types | PANTHER Pathway DB and Biomodels repository ChEBI, Gene/protein DBs (e.g. Uniprot), GO, SBO | 15901 entities, 12125 events, 571 modifications |
| Gene Regulation Ontology (GRO) [35] | 300 abstracts | human gene regulation and transcription | 174 entity types, 126 event types | Gene Regulation Ontology | 11819 entities, 4837 events, 5241 event instances, 3832 relations |
| Gene Regulation Network (GRN) [24] | 201 sentences (from abstracts) | sporulation in *Bacillus subtilis* | 12 entity types, 9 event/relation types, 6 interaction types | Bibliome team, LIPN | 917 entities, 495 biochemical events, 334 interactions |
| Bacteria Biotopes (BB) [23] | 131 documents (web pages & encyclopedia articles) | microbial habitats | 9 entity types, 2 event types | OntoBiotope-Habitat ontology | 5183 entities, 2260 relations |

The following sentence illustrates the event extraction process: “TGF-beta mediates RUNX induction, and FOXP3 is efficiently up-regulated by RUNX1 and RUNX3” (Fig. 1). First, in the entity recognition step, TGF-beta, RUNX, FOXP3, RUNX1 and RUNX3 are identified and annotated as protein mentions. Next, trigger words indicating the presence of events are identified, and events are detected and labeled as edges. In this example, one event, gene expression, is indicated by the nominalized verb ‘induction’ whose theme is RUNX, a protein. The second event, positive regulation, is indicated by the verb ‘mediates’ whose cause is TGF-beta, a protein, and whose theme is the gene expression event. The third event, positive regulation, is indicated by the verb ‘up-regulated’, whose causes are two proteins RUNX1 and RUNX3, and the theme is another protein FOXP3. The ‘up-regulated’ token produces positive regulation event with overlapping edges, and the resulting structure needs to be pulled apart to construct a valid event.

**Figure 1.**
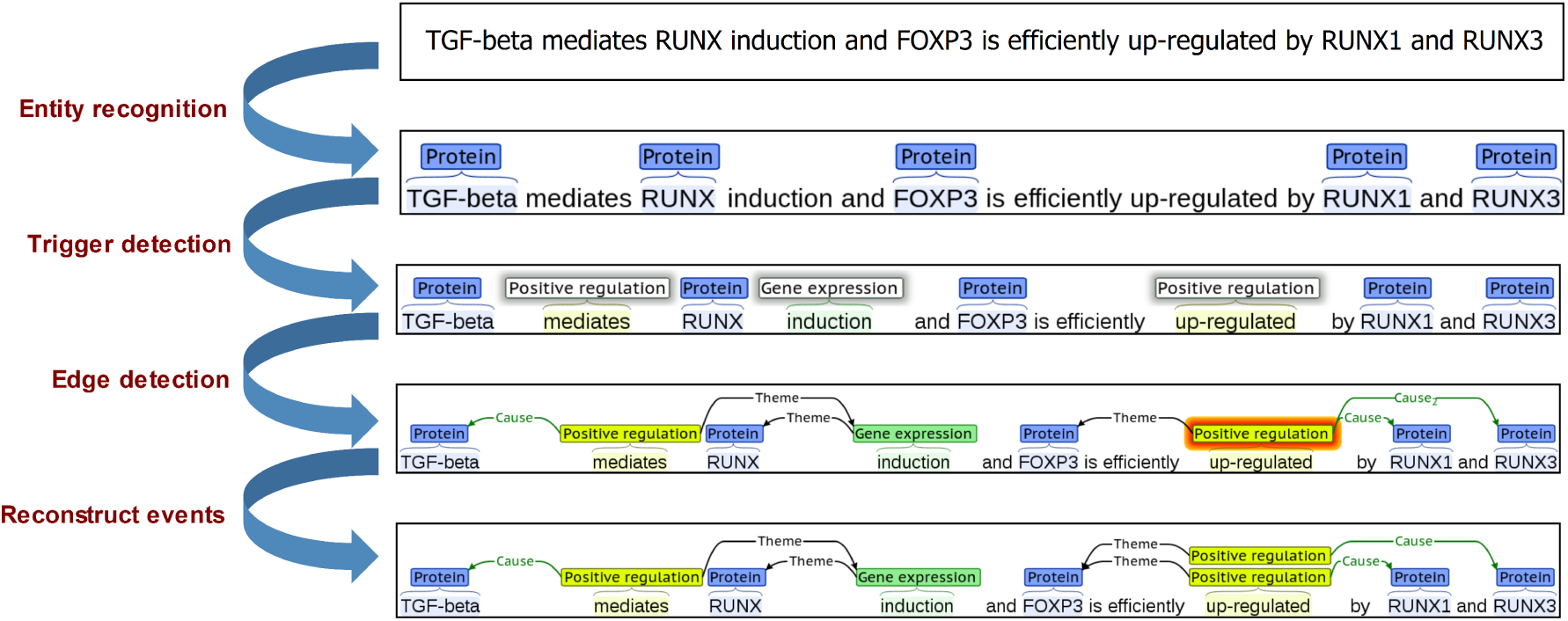
Example of event extraction. Entities and event triggers are shown with their types above their corresponding text and event participants as arcs marked with roles. Each type is shown in a di_erent colour, as indicated.

Although the figure depicts the detection of events based on pipeline approaches (see “Event Extraction Methodologies” section), it nevertheless gives a quick guide on the general procedure of event extraction for illustration purposes. As demonstrated in the example above, event extraction depends heavily on the domain-specific concepts and ontologies. Henceforth, the development of biomedical NLP application has called for a community-wide effort to establish and standardize these ontologies (and corpora). In this review we focus our discussion on the BioNLP Shared Task (BioNLP-ST) series, the first community-based effort to target the fine-grained extraction of bio-events in biological text [19, 20, 21].

## BioNLP Shared Tasks

To promote the development of NLP applications dedicated to the biomedical domain, a series of biennial competitions are being held under the title BioNLP-ST. At each round, BioNLP-ST organizers establish protocols for the competition, provide common resources comprising a rich collection of knowledge bases and annotated training corpora, and set evaluation benchmarks to measure and rank the performance of competing tools. Such a structured effort is key to creating a common framework that describes the needs of biologists, and to guiding development of cutting-edge BioNLP applications with measurable performance.

The BioNLP-ST series was initiated in 2009 with an event extraction challenge using the Genia corpus [22], and has since evolved substantially through the addition of new subtasks; expansion of its scope to extracting events from full papers in addition to abstracts; and, growing sophistication of its resources. The most recent BioNLP in 2013 [21] included six STs which are summarized in Table 1.

Further, BioNLP-ST competitions encourage participants to deposit code and trained models in public repositories; introduce new approaches that decrease reliance on deeply-annotated data; and, improve database curation. Post-competition, many participants continue to improve their methods, providing continued state-of-the-art performance.

### Evaluation Metrics

The performance of a BioNLP system is typically measured based on its true and false positive observations, which are used to calculate ***precision, recall*** and ***F-score*** metrics. Precision describes the fraction of retrieved documents or extracted events that are relevant, as deemed by the ground truth. Recall (also called sensitivity) measures the fraction of relevant documents or entities that are retrieved by a method. The F-score measure is the harmonic mean of precision and recall. In most cases, precision and recall are inversely related: precision scores usually decrease as the retrieved documents are increased at the cost of the recall, and vice versa. The F-score measures the trade-off between precision versus recall. For additional measures, such as ‘receiver operating characteristics’ (ROC) curves, ‘area under the ROC curve’ (AUC), ‘slot error rate’ (SER) and others, see [23, 24].

### Corpora

Table 1 provides descriptive statistics for several BioNLP-ST corpora mentioned in this paper, many of which follow event annotation guidelines that were introduced by the Genia event corpus. Additional collections of corpora are available on the web, notably under the ‘Resources’ sections at the BioCreAtivE [25] and NaCTeM [26] websites. The ‘Collaborative Annotation of a Large Biomedical Corpus’ (CALBC) project offer a list of their gold standard corpora [27]. Hakenberg at the Icahn School of Medicine at Mount Sinai Hospital provides a reasonably comprehensive and well-organized list [28]. Further, the Institut für Informatik at Humboldt-Universität zu Berlin provides a large collection covering protein-protein, drug-drug and disease-treatment interactions, gene variations and chemical compounds - many of which have been released in the standardized BioC format [29].

## Event Extraction Methodologies

Most event extraction analysis methods can be categorized as pipelines, which process texts in a cascading manner, where the output of the preceding step is propagated as input to the next. The event extraction pipelines in the BioNLP-ST competitions can be classified into (i) statistical; (ii) symbolic; or (iii) hybrid methods. On the other hand, statistical joint inference methods offer an alternative to this pipeline architecture, as they usually perform their analyses in one phase.

### Pipeline-based statistical approaches

In BioNLP, statistical pipelines usually employ a supervised learning framework, using a provided training set of manually annotated text. The methods sequentially deploy modules comprising event triggers, arguments, etc., to tune model parameters, such that a defined fitness function is optimized on the training set. The trained models are then applied to a test set (unannotated and hitherto unseen).

*TEES* is an open source event extraction system developed at the University of Turku [36]. It considers events as text-bound graphical structures, where the nodes of the graph consist of named entities and event triggers, and the edges represent relations and event arguments [37]. Within this graphical representation, event extraction is formulated as the task of detecting the nodes and edges. TEES breaks down these tasks into a series of multi-class classification problems. It uses a support vector machine (SVM) [38] to extract an event graph from the parsed text in a four-stage pipeline [37]. The *trigger detection* stage of the pipeline predicts event classes for each non-named entity token, while the *edge detection* stage identifies corresponding arguments of the detected events through a multi-class SVM that assigns candidate edges to one of several semantic roles, such as theme or cause. The constraints over argument roles are learned automatically from the corpus, thus allowing the system to be automatically adapted to new corpora. The *unmerging* stage converts overlapping events into structurally valid ones, while the *modality detection* stage, an optional procedure, may be used to classify trigger nodes as negated or speculated events.

TEES uses several feature sets for trigger and edge detection, including dependency-based features derived from a full syntactic analysis of the sentence. For this, dependency trees are generated by converting the phrase-structure output produced by the McClosky-Charniak parser [39] into the Stanford *collapsed dependencies with propagation* format [40, 37], which is a compact representation of a dependency graph that does not necessarily represent every word in the sentence.

TEES has been a strong contender in BioNLP-ST competitions. In 2009, it placed first in the GE task with an F-score of 51.95% [19]. In 2011, it placed third in the GE task behind Riedel’s joint model (discussed in “Statistical joint inference models” section), and first in four of the remaining seven tasks [31, 32]. In 2013, TEES again performed strongly, placing first in the CG, GRO and BB (tasks 2 and 3) and second in the GE and PC tasks, behind EventMine [36]. It also performed well in the DDIExtraction task (concerning the recognition of drugs and extraction of drug-drug interactions that appear in biomedical literature), participating in both 2011 and 2013 [41, 42].

TEES has been employed in large-scale text mining applications, for example in extracting events from 19 million unannotated PubMed abstracts [43], and in generating the EVEX dataset, a collection of over 40 million biomolecular events extracted from all PubMed abstracts and PubMed Central full text articles [44]. [8] proposed a modified version of TEES augmented with features derived from the EVEX dataset. This resulted in a modest improvement over TEES, leading to a first place in the GE task of the 2013 BioNLP-ST competition, with an F-score of 50.97%. Other applications of TEES include its use in: the CellFinder literature curation pipeline [45], the BioContext integrated text mining system [46], and semi-automated knowledge extraction workflow [47]. With growing interest towards adopting such open source tools as modules in larger BioNLP systems, TEES is a leading event extraction system that can be further improved on by incorporation of more advanced coreference resolution techniques, and addition of methods for adaptive learning from multiple corpora to detect events spanning multiple sentences from heterogeneous texts.

*EventMine* is an event extraction system developed by the National Centre for Text Mining (NaCTeM) [26, 48]. Similar to TEES, EventMine employs a modular architecture: a *trigger/entity detector* to find and classify word candidates (e.g., as ‘binding’ or ‘entity’); an *argument detector* to identify and categorize candidate relations between detected triggers and entities according to semantic roles; a *multiple argument detector* to reconstruct event structure properly by considering all possible combinations of detected relations in the previous step, and to categorize these events into event classes; and a *hedge detector* to characterize the modality of detected events as being speculated or negated. EventMine, as in TEES, applies more efficient hyper-parameter tuning method to automatically adapt with different tasks and it is stored in a global configuration file. Nonetheless, manual inspection is only required when users change the task-specific settings and other options which are outlined in [49].

In each module, the corresponding model solves a multi-class multi-label classification problem with a one-versus-rest SVM classification scheme [50]. EventMine uses a rich set of features for the classification problem [17], including token-based features, neighbouring word features, and features that are generated by GDep [51] and Enju [52] parsers. In contrast to TEES, features generated in EventMine are mapped to a hash table to reduce the memory usage. Additionally, the system introduces a rule-based coreference resolution, which further boosts its performance [53].

In the 2013 BioNLP-ST competition, EventMine ranked first and second in the PC and CG tasks with F-scores of 52.84% and 52.09%, respectively [48, 6]. Similar to the combined TEES + EVEX system [8], EventMine has been used to improve event extraction across multiple corpora with partially overlapping semantic annotations, such as the GE and ID corpora [17], with the aim of reducing the need for manual annotation of existing semantic types in new corpora and improving learning on heterogenous texts [17, 53, 5].

Further, EventMine has been extended to EventMine-MK [54], which uses five dimensions of textual context surrounding events to enhance its trigger detector module. These five dimensions consist of the following. (i) *Knowledge type:* the general information expressed by the event; (ii) *Certainty level:* the confidence of an event being expressed; (iii) *Polarity:* the truth conditions of an event; (iv) *Manner:* the intensity level of an event; and (v) *Source:* whether an event occurred in the current paper or a previous study. EventMine-MK has been evaluated on extracting negated and speculated events, a subtask of the 2009 BioNLP-ST competition. With the integration of meta-knowledge, EventMine-MK outperformed the other state-of-the-art systems that participated in this task [54].

*StanfordEE*, Stanford University’s entry to the 2009 BioNLP-ST competition [55] introduced a re-ranking dependency parser that offered an alternative representation of the event structure via a *transformation and trigger detector*. In this system, the candidate events for each sentence are transferred into a dependency tree structure. The nodes in this tree comprise entity mentions, triggers, and a virtual ‘root’ node. Then an *edge detector* predicts links between tree nodes, and a *conversion unit* transforms the dependency trees back to their pseudo-syntactic event structures. Finally an *event structure selector* picks the best candidate of pseudo-syntactic event structure using multiple decoders from the MSTParser [56], which produces both non-projective and n-best parses.

For detecting events, a logistic regression classifier with L2 regularization is employed to label individual token independently based on several features including surface, syntactical, contextual, bag-of-word counts, and domain specific features such as ontology classes.

In the 2011 BioNLP-ST competition, StanfordEE achieved 3rd, 5th, and 7th places in the ID, EPI and GE tasks with F-scores of 50.60%, 31.20%, and 50.00%, respectively [55]. In 2011 it also competed in conjunction with the UMass joint event extractor [57], as described in “Statistical joint inference models” section.

The StanfordEE stands out with its following features: First, the parsing of event structures can be applied over full text instead of individual sentences, allowing events to be modeled as discourse patterns over documents. Second, the system is agnostic to the parsing model. Third, it is flexible in its re-ranker module, which can be combined with multiple parsers and multiple trigger recognizers. Fourth, it employs domain independent features for detecting event anchors, with the exception of conversion of event structures in annotated ST corpora to tree dependency structures. However, in practice, representation of event structures as trees is frequently violated. In such cases, it is not clear how the system can be applied for downstream analyses, such as resolving anaphoric constructs.

*General Remarks:* While many state-of-the-art statistical systems employ a ‘pipeline’ architecture, a drawback of this approach is the cascading of the errors that may be introduced in the early stages of the process. For example, if a trigger is missed by a preceding module, it would not be recovered in the next stage. Furthermore, during learning parameters in the edge detection phase each identified trigger token will predict at least one argument, which might not establish a true event type. This will increase the false positive rate, and decrease the false negative rate. To maintain a balance between precision and recall, one could instead pass several token candidates to the next stage, and subsequently eliminate triggerless events to reduce false positives. Such a strategy is employed by EventMine, targeting high recall in its four event extraction detection modules [53, 17] by weighing positive instances in combination with trigger filtering techniques and hand crafted rules during the edge detecting phase.

Another significant limitation of the pipeline-based statistical approaches is the use of local classifiers (those trained without the knowledge of any global constraints). In contrast to the joint inference approaches described in “Statistical joint inference models” section, local classifiers predict event types independently from arguments and their roles, confining the global correlation between these variables.

Overfitting is a general problem among the statistical models discussed in this section. This refers to tuning a system to fit the trained data highly accurately, while making poor predictions on the test data. It happens when the model ‘learns’ the nonexistent ‘structure’ in the random noise, rather than tuning to the desired features in the training set. While cross-validation techniques are widely applied to minimize overfitting, sparsity of a feature in the training set makes this challenging [58]. Generalizing multiple features into fewer categories may alleviate the latter problem to a certain extent, yet it may also introduce over-generalized features, which, with enough training instances, may increase false positive rates. Furthermore, while some feature sets may provide good prediction with particular data, others may not [17].

Despite these drawbacks, the pipeline-based statistical approaches are consistently the top performers in BioNLP-ST competitions. They formulate tractable problems even for large datasets, extract salient information, and provide verbose outputs that can be used to refine these algorithms.

### Pipeline-based symbolic approaches

A good alternative to event extraction using statistical approaches is employing symbolic approaches, especially when the underlying text is domain specific. Such pipelines extract events and their arguments based on set rules that are either manually defined, or automatically learned from training data. There were several pipeline-based symbolic approaches that participated in the BioNLP-ST events, some of which displayed highly competitive performance.

*BioSem* is a symbolic event extraction model that has two phases: a learning phase and an extraction phase [59]. During the learning phase it automatically constructs a dictionary using event triggers that are derived from the training data. It then generates lexical patterns from each event by matching events with syntactic layer containers (e.g., chunk, phrase or clause) that have event triggers and their arguments. If an event is matched with a container, it generates a pattern by extracting a list from a pre-defined feature set associated with that container. During the extraction phase, it uses the obtained dictionary and patterns to call events in an input text. In 2013 BioNLP-ST competition BioSem placed third in GE task 1, with an F-score of 50.68% [59]. Recently, the system has been extended with SVM into a hybrid pipeline, and specializes in extracting drug-drug interactions [60].

*OpenDMAP* pipeline combines elements of several community based ontologies in defining its internal structure that contains defined classes and slots, and then uses the resulting ontology to validate semantic constraints during event extraction [61]. Doing so, it provides complete relational structures in one step, defining the linguistic context of an event predicate along with its arguments. For trigger detection, OpenDMAP employs a dictionary lookup using ConceptMapper [61]. The system’s reliance on hand crafted rules, while highly precise, resulted in comparatively poor recall scores during the 2009 BioNLP-ST competition. Overall the tool achieved an F-score of 13.45% on task 1 and 22.33% on task 2.

*ConcordU*. A different strategy to event extraction was proposed by the Concordia University team [62]. The ConcordU system treats biological event extraction as a discourse interpretation across sentences. This is achieved in a two-phase approach: syntax-driven composition phase and mapping phase. During syntax-driven composition phase, a general semantic representation is crafted from syntactic analysis of texts. This involves (i) a dictionary lookup; (ii) a statistical measurement of trigger expressions including trigger polarities; and (iii) execution of the BLLIP reranking parser [39]. During the mapping phase, it maps relevant parts of the semantic representation to event instances according to the event definitions and constraints. During the 2009 BioNLP-ST competition, the ConcordU system placed third in the GE task 1 and first in GE task 3 [63]. It also performed moderately-well in 2011 BioNLP-ST [62]. We note that, unlike OpenDMAP and BioSem, the ConcordU system included argument identification rules to address coreference resolution.

*General Remarks:* Results from the BioNLPST competitions demonstrate that pipeline-based approaches generally attain high precision but low recall. The low recall can be explained by the reliance of these systems on precisely defined triggers and entity mentions to extract events: if a particular rule is too restrictive, then the procedure used to recover the event will fail, yielding low recall. Furthermore, many symbolic systems do not make use of the full syntactic structure of the text, limiting accurate extraction of discourse-level elements such as negation, speculation, nominalization, coordination, anaphora and coreference [64].

However, symbolic approaches do offer several benefits over statistical approaches. Symbolic approaches flexibly combine input from heterogeneous systems, which is very difficult to achieve using statistical methods that require fixed feature collections. Although the feature space could be extended in statistical approaches, it is very difficult to understand which parts of the model are significant, without exhaustive feature testing. They are not trained on a model; instead they use hand-crafted rules that are easier for humans to comprehend than the patterns computationally learned in statistical methods. In this regard pipeline-based symbolic systems are easier to maintain, troubleshoot, improve, and adapt to other domains. Finally, in some cases, symbolic approaches may capture patterns that are not well handled by statistical methods.

DRUM (Deep Reader for Understanding Mechanisms) is a recent symbolic-based event extraction system, more precisely cognitive point of view, which combines syntactic structure with full fledged comprehensive logical understanding of text [65]. While this model represents a leading-edge effort for symbolic-based model to understanding big and complex mechanisms, we exclude as our preferred symbolic approach because DRUM did not report results over the BioNLP-ST datasets. We believe that substantial experiments are required to proof its effectiveness on event extraction datasets.

### Hybrid Pipelines

Hybrid approaches combine the strengths of the previous two approaches to keep the high recall rates of the statistical systems, while providing high precision of the symbolic systems.

*NCBI-ASM*. National Center for Biotechnology Information (NCBI) developed a system based on approximate subgraph matching with three modules: (i) rule induction, (ii) sentence matching, and (iii) rule set optimization to increase the overall precision [66]. The system uses an all-paths graph kernel [67] among event arguments to extract events, and a distributional similarity model. The NCBI-ASM approach significantly increases the extraction precision at a high level. In the 2013 BioNLP-ST competition, this system achieved moderate F-scores of 46.38% and 48.93% (3rd and 4th places) in the CG and GE Tasks, respectively.

*TEES+GETM*. Van et al. [68] developed a hybrid framework combining the trigger detection unit of TEES with GETM (the Ghent Text Mining relation detection framework) to produce high recall rates, while benefiting from the relatively high precision rates inherited from both systems. However, this combination results in many false positives, mainly due to the differences between the two systems in building features. GETM extracts entity relations between gene symbols and domain terms, while TEES is a generalized biomedical event extraction tool.

*General Remarks:* By adopting hybrid-based frameworks, the compound benefits can be absorbed from rules, graphs and statistical approaches in contrast to applying each of these methods individually. However, the displayed performance of such tools indicate that integration of methods across paradigms is a nontrivial task, and would require further research to deliver the potential of the approach.

### Statistical joint inference models

Almost all the systems discussed above make a strosng independence assumption between trigger words and event arguments, by treating their detections as separate tasks. In doing so, these systems essentially ignore any information provided by the downstream modules, leaving these systems susceptible to cascading errors. Hence, methods that can jointly infer the entire event structure would have distinct advantages over the methods that assume these tasks to be independent classification problems. Joint inference methods can make use of dependencies between tasks, for instance capturing correlations between the evidence for possible arguments, and the detection of event triggers.

*Markov Logic Networks (MLNs)* provide a framework for statistical relational learning, an emerging area of research concerned with reasoning and machine learning in domains that are characterized by uncertainty and complex relational structure [69]. Markov logic is a probabilistic extension of first order logic, and incorporates ideas from probabilistic graphical models [70]. The MLN syntax uses sets of predicates, hidden and observable, and weighted logical formulae to encode constraints about a system. Intuitively, the most probable systems are those that violate the fewest constraints, where violating a formula with a higher weight incurs a large penalty [69].

In this approach, ‘hidden’ predicates represent the predicted targets of the system. Examples of query predicates include *event(i)*, which is true if token at index i is a clue word, or *eventType(i*, e), which is true if an event at token index *i* has event type *e*. Observable predicates capture properties of the input sentence, such as “token *i* has word *w*”, or “token pair *(i,j)* has dependency label p”. On the other hand, the logical formulae comprise both local and global sets. Local formulae connects the observable predicates to the hidden predicates. Global formulae capture constraints or correlations between hidden predicates, and are one of the main advantages of MLN over conventional classifiers. These formulae can capture hard constraints that enforce the validity of events, such as structural constraints like “all events have a type”, and are learned deterministically. Softer correlations can also be included formulaically to improve the performance of the system, such as formulae that capture linguistic regularities.

Riedel et al. [71] modeled event extraction as a link-prediction problem [72] centered around MLN method, where the existence of relationships or links between entities are inferred from direct and indirect evidence. In this case, the links in question are argument structures that exist between tokens in a given sentence. For example, when the first and third tokens in a sentence are labeled as triggers of regulation events, a system might predict a link between these two tokens. In this model, events and their associated arguments are converted into sets of labeled clue words and token-to-token links. In the 2009 BioNLP-ST competition, this system ranked fourth in task 1 with an F-score of 44.35% and first in task 2 with an F-score of 43.12%. However, it performed poorly in detecting binding events. The reasons could be attributed to: incorrect heuristics when converting detected links into the corresponding event structures; inconsistencies in the training data; and, the performance of the learning algorithm used for training the model [71].

While the previous systems treat argument detection as a token classification problem, Poon at al. [73] proposed an approach that analyzes edges predicted by the Stanford dependency parser [40] to mark edges that are in argument paths. Their system uses a number of heuristic methods to fix common errors resulting from syntactic parsing, and allow the model to learn and exploit correlations between sequences of dependent edges. The model is an extension to the open source Alchemy software package [74], and makes use of parallel processing for learning and inference steps to achieve high computational efficiency. Although the team did not participate in the BioNLP-ST competitions, it was able to achieve an F-score of 50.0% on the 2009 BioNLP-ST competition test set.

More recently, Venugopal at al. [75] developed a method for incorporating high dimensional features into MLN, overcoming some of the associated computational bottlenecks that had forced previous Markov logic approaches to use simpler feature sets. This method learns high-dimensional features using SVMs, and encodes the resulting output as low-dimensional constraints in MLN. To further reduce the computational demands of inference in MLN, the network is partitioned into several disconnected components and solved independently. This approach led to the second-best F-score on the 2009 GE dataset of BioNLP-ST competition (58.16%), and the highest F-scores on the 2011 and 2013 on GE task 1 of BioNLP-ST competitions (58.07% and 53.61%, respectively).

The MLN based event extraction methods posses several remarkable properties. First, MLNs allow the capture of global correlations between variables and decisions, such as the constraint that a predicate can only have one argument. Second, they allow a ‘divide and conquer’ approach, where inference can be performed on the subgraphs of a partitioned network without having to fully instantiate the complete graph [76]. Finally, MLN models implement a single stage operation, as opposed to an analysis pipeline, substantially simplifying software development, implementation and management [71, 77].

*Dual and Stack Decomposition Models*, like MLN models, also offer a ‘divide and conquer’ strategy [78]. They allow reformulation of complex inference problems into efficiently solvable subproblems. These subproblems are linked together with linear constraints, which enforce agreement between their solutions. The agreement constraints are integrated into an optimization problem using Lagrange multipliers, and an iterative algorithm is used to minimize the resulting cost function. The subproblems are chosen in such a way that they can be solved efficiently using exact combinatorial algorithms [78].

In this group, the UMass [57] is a dual decomposition system comprising three joint models. Its first model provides joint trigger and argument extraction using a simple, efficient and exact inference algorithm; its second model ensures consistency between events in hierarchical regulation structures; while a third model combines the first two models to ensure consistency between arguments of binding events. In the 2011 BioNLP-ST competition, the UMass system placed first in the GE task 1 on the full papers collection, with an F-score of 53.14% [79]. The UMass system provides significant improvements over competing joint approaches, and is robust across multiple corpora. However, it is neither modular nor flexible, requiring extensive modifications in its learning and inference components to adapt to new domains and to changes in the corpora.

Another model that breaks down a process into multiple subprocess is the FAUST system [80]. FAUST uses a stacked framework employing the UMass dual decomposition model [57] and the StanfordEE [55], where predictions from the Stanford parser serve as features for the UMass system. This combination of the two competitive models outperforms the UMass system, with substantially improved accuracy. In the 2011 BioNLP-ST competition, FAUST obtained first place in three out of four tasks: first place in GE tasks 1 and 2 and the ID track with F-scores of 56.0%, 53.9% and 55.6%, respectively, and second place in the EPI track with an F-score of 35.0%.

*Other Joint-based Models*. RecUrsive Pairwise Event Extractor (RUPEE) [81] detects detects pairwise interactions between text entities recursively in two steps: (trigger, theme) pair extraction that identifies the trigger words with one of their arguments; and postprocessing that adds extra arguments to binding and regulation events. While the former step uses a multiclass classifier based on SVM, one for each event class, the later one adopts a binary SVM to classify: optional cause arguments for regulation events, and possible multiple arguments for binding events. RUPEE did not participate in the BioNLP competitions, however, it outperformed the previous joint models for 2013 GE task 1 and achieves third for 2011 GE task 1 of the BioNLP-ST datasets with F-scores of 54.4% and 55.6%, respectively.

SEARN is an algorithm for integrating SEARCH and LEARNING to solve structured prediction problem, such as events. Vlachos et al. [82] formulates the problem for events by representing each instance/sentence as a sequence of multiclass predictions through defined hypotheses. In each iteration, a cost-sensitive classification (CSC) is generated for each action and for each instance; at the end, the learned hypotheses are used for predicting events and their arguments. The intuition behind SEARN is that good performance on the derived classification problems implies good performance on the structured prediction problem. SEARN’s high modeling flexibility over dual decomposition methods and MLN systems did not guarantee to provide a good performance in BioNLP-ST 2011.

HDS4NLP [83] decomposes the overall task into a multi-class problem of classifying (trigger, argument) pairs using SVM. The HDS4NLP system ranked 6th on the GE task with an F-score of 43.03%; however, post competition it outperformed the winner of this task after fixing a software bug, achieving an F-score of 51.15%.

Kordjamshidi et al. [84] proposed another joint based model customized specifically for Bacteria Biotopes (BB) task based on the framework of structured output prediction [85], similar to SEARN. The model, ‘Spatial Role Labeling (SpRL),’ uses a large collection of local and global (i.e. document level) features of words, phrases, sentences and paragraphs for extracting entities and relations. In addition, it relies heavily on the OntoBiotope [86] ontology of microbe habitats and the NCBI bacterium taxonomy database [87] to increase the accuracy of detecting entities. SpRL is trained on structured SVM (SSVM) using link-and-label (LAL) framework [88]. Empirical evaluation of this model conducted solely on BB dataset of the BioNLP-ST 2013 and reported the best F-score measure of 0.22%. However, the system did not generalize to other shared tasks datasets.

*General Remarks:* As discussed, sequential pipelines with multiple stages potentially suffer from error propagation, since event triggers and arguments are predicted in isolation by independent local classifiers. The joint inference models were developed, in particular, to address this issue. A shared characteristic of joint models is the implementation of inference over complete event structures for a given sentence. By allowing information to propagate across events and arguments, joint inference can facilitate mutual disambiguation, potentially leading to substantial gains in predictive accuracy. However, because they address the entire event extraction problem globally, joint inference models are more difficult to build. Also, popular implementations of joint inference methods are computationally costly [57, 76], notwithstanding their suitability for ‘divide and conquer’ strategies, potentially allowing for parallel processing to reconstructing events over large scale text and to minimize latency in the execution time.

## Discussion and Conclusion

While state-of-the-art event extraction methods hover around 50% accuracy on average over multiple shared tasks (F-scores), they are finding use in several real-world applications [89, 90]. For the event extraction technologies to find a wider uptake, several challenges need to be addressed.

**1**. Parsers utilized in these systems affect the overall performance of event extraction models because they are trained on different corpora, each characterized by their own sets of annotations and syntactic structures. Although combination models trained on different domains outperform a single baseline models as reported in [91], the formats generated by these parsers remain limited; for example, some representations are better at detecting regulation events than binding events [92].

**2**. The performance of event extraction systems over full text analysis is highly dependent on the correct identification of biomedical entities, such as gene and protein mentions. Defining the textual boundaries that encapsulate an entity is a complex procedure, as entities in the biomedical literature could have multiple synonyms, often with different acronyms and abbreviations [93]. Furthermore, disambiguating entity mentions with respect to other entities is perhaps the hardest part of NER; for example, in the phrase “left breast cancer” it is not clear whether the author intended to refer to cancer in the left breast, or a type of cancer (however unlikely, biologically) that is specific to the left breast [1].

**3**. While they are commonly adopted for event extraction systems in BioNLP-ST competitions, ontologies are primarily developed for the needs of biologists, not text mining. The conceptual model defined by an ontology and how these concepts appear in text are fundamentally different [94] from those required in BioNLP. Lexical resources that provide intermediate knowledge are required to bridge ontologies and biomedical text, that is the gap between ontology design and the requirements of the BioNLP community [95].

**4**. Corpora employed to train event extraction systems tend to perform best when information is stated explicitly and locally in text. When statements are not clearly specified the correct meaning must be inferred from context - such as previously stated information, or the background knowledge of the reader. For example, Buyko et al. [96] suggest that their weak classification performance of gene regulation relations was partly because their system was incapable of making use of inferential knowledge. In addition, the level of semantic detail required for annotating corpora is a crucial factor impacting the performance of event extraction systems.

The scheme used for annotating a corpus generally consists of a subset of concepts from a reference ontology [35]. For example, the event ontology embedded in the GENIA corpus is based on a subset of 36 classes that are defined in GO, mostly using general concepts in the upper levels of the concept-hierarchy. While using a broad concept to annotate a large range of related ideas will likely increase the coverage of annotations, it nonetheless may also introduce problems. For example, the regulation events in GENIA are used to express general causal relationships in addition to biological regulatory processes [16]. An intermediate solution would be to annotate each corpora with low level of detail and then combining them into a single resource; however, the differences in semantic scope, compounded with dissimilar annotation formats and guidelines, might inhibit such integration [97]. Another possible solution would be to annotate a corpus with handful amount of semantic detail that overlaps with multiple corpora, such as BM corpus (Big Mechanism project [9]). Given that annotating a corpus is time and cost consuming, semisupervised learning approaches or active learning approaches might be employed [2].

**5**. Coreference resolution, which finds all expressions that refer to the same entity in a text, is poorly resolved in event extraction systems [1]. Recent research [98] confirms that co-reference resolution is key to accurate relation detection. This topic remains an active and open area of research.

**6**. Event extraction, currently implemented as a supervised learning task, relies heavily on the annotation scheme of the training data, and thus may drop significantly in performance when the schema changes, even within the same domain [99]. One of the problems usually encountered with supervised models is that they are specifically tailored to features of the corpus upon which they are trained. Consequently, their functionality is limited to the semantic types that they are trained to identify. Domain adaptation approaches at this stage filter out redundant and contradicting labeling across different corpora, and then merge the corpora in order to train a single combined model.

Instead of adapting corpus annotations to specific tasks, an alternative approach is to develop and train unsupervised models, for the creation of larger, more flexible corpora. Another option would be to take advantage of increases in the computational power, algorithms and data, allowing models to be learned automatically through deep processing [100]. A careful construction of representational models that utilize the characteristics of words around the context into event extraction could achieve a comparable result as reported in the recent study [101].

Other challenges, including detection of negation and speculation, normalization of relations (e.g. gene names, abbreviations, etc.), inclusion of non-textual material in biomedical event extraction pipelines, and the automatic integration of event extraction models with biological pathways (facilitate the understanding of biological interactions) [102, 2, 54].

The paradigms explored in this paper have strengths and weaknesses. While building on these strengths in hybrid or stacked systems is an appealing option, integrating them is a challenging task. Resolving the challenges facing event extraction methods would provide additional benefit to the biomedical community, for example allowing development of improved event-based information retrieval systems, the construction of knowledge bases, and enabling us to uncover latent knowledge aggregated in the published literature and to formulate new hypotheses.

## Acknowledgment

We thank the Sequencing Lab and the Bioinformatics Technology Lab (BTL) at Genome Sciences Centre, British Columbia Cancer Agency for their assistance with this article.

## References

1 Rebholz-Schuhmann, D., Oellrich, A., Hoehndorf, R.: Text-mining solutions for biomedical research: enabling integrative biology. Nature Reviews Genetics 13(12), 829–839 (2012)

2 Zhou, D., Zhong, D., He, Y.: Biomedical relation extraction: From binary to complex. Computational and mathematical methods in medicine 2014, 18 (2014)

3 Ananiadou, S., Thompson, P., Nawaz, R., McNaught, J., Kell, D.B.: Event-based text mining for biology and functional genomics. Briefings in Functional Genomics, 015 (2014)

4 Li, C., Liakata, M., Rebholz-Schuhmann, D.: Biological network extraction from scientific literature: state of the art and challenges. Briefings in bioinformatics, 006 (2013)

5 Pyysalo, S., Ohta, T., Miwa, M., Cho, H.-C., Tsujii, J., Ananiadou, S.: Event extraction across multiple levels of biological organization. Bioinformatics 28(18), 575–581 (2012)

6 Ananiadou, S., Pyysalo, S., Tsujii, J., Kell, D.B.: Event extraction for systems biology by text mining the literature. Trends in biotechnology 28(7), 381–390 (2010)

7 Miwa, M., Ohta, T., Rak, R., Rowley, A., Kell, D.B., Pyysalo, S., Ananiadou, S.: A method for integrating and ranking the evidence for biochemical pathways by mining reactions from text. Bioinformatics 29(13), 44–52 (2013)

8 Hakala, K., Van Landeghem, S., Salakoski, T., Van de Peer, Y., Ginter, F.: Evex in st’13: Application of a large-scale text mining resource to event extraction and network construction. ACL 2013, 26 (2013)

9 Cohen, P.R.: Darpa’s big mechanism program. Physical Biology 12(4), 045008 (2015)

10 Buyko, E., Faessler, E., Wermter, J., Hahn, U.: Event extraction from trimmed dependency graphs. In: Proceedings of the Workshop on Current Trends in BioNLP: Shared Task, pp. 19–27 (2009). Association for Computational Linguistics

11 Navlakha, S., Kingsford, C.: The power of protein interaction networks for associating genes with diseases. Bioinformatics 26(8), 1057–1063 (2010)

12 Ozgur, A., Vu, T., Erkan, G., Radev, D.R.: Identifying gene-disease associations using centrality on a literature mined gene-interaction network. Bioinformatics 24(13), 277–285 (2008)

13 Tatonetti, N.P., Patrick, P.Y., Daneshjou, R., Altman, R.B.: Data-driven prediction of drug effects and interactions. Science translational medicine 4(125), 125–3112531 (2012)

14 Tari, L., Anwar, S., Liang, S., Cai, J., Baral, C.: Discovering drug-drug interactions: a text-mining and reasoning approach based on properties of drug metabolism. Bioinformatics 26(18), 547–553 (2010)

15 Miwa, M., Sætre, R., Kim, J.-D., Tsujii, J.: Event extraction with complex event classification using rich features. Journal of bioinformatics and computational biology 8(01), 131–146 (2010)

16 Kim, J.-D., Ohta, T., Tsujii, J.: Corpus annotation for mining biomedical events from literature. BMC bioinformatics 9(1), 10 (2008)

17 Miwa, M., Pyysalo, S., Ohta, T., Ananiadou, S.: Wide coverage biomedical event extraction using multiple partially overlapping corpora. BMC Bioinformatics 14(1), 175 (2013)

18 Kim, J.-D., Wang, Y., Yasunori, Y.: The genia event extraction shared task, 2013 edition-overview. ACL 2013, 8 (2013)

19 Kim, J.-D., Ohta, T., Pyysalo, S., Kano, Y., Tsujii, J.: Overview of bionlp’09 shared task on event extraction. In: Proceedings of the Workshop on Current Trends in BioNLP: Shared Task, pp. 1–9 (2009). Association for Computational Linguistics

20 Kim, J.-D., Pyysalo, S., Ohta, T., Bossy, R., Nguyen, N., Tsujii, J.: Overview of bionlp shared task 2011. In: Proceedings of the BioNLP Shared Task 2011 Workshop. BioNLP Shared Task ‘11, pp. 1–6 (2011). Association for Computational Linguistics

21 Nédellec, C., Bossy, R., Kim, J.-D., Kim, J.-j., Ohta, T., Pyysalo, S., Zweigenbaum, P.: Overview of bionlp shared task 2013. In: Proceedings of the BioNLP Shared Task 2013 Workshop. BioNLP Shared Task ‘13, pp. 1–7 (2013). Association for Computational Linguistics

22 Kim, J.-D., Ohta, T., Tateisi, Y., Tsujii, J.: Genia corpus a semantically annotated corpus for bio-textmining. Bioinformatics 19(suppl 1), 180–182 (2003)

23 Bossy, R., Golik, W., Ratkovic, Z., Bessières, P., Nédellec, C.: Bionlp shared task 2013–an overview of the bacteria biotope task. ACL 2013, 161 (2013)

24 Bossy, R., Bessières, P., Nédellec, C.: Bionlp shared task 2013-an overview of the genic regulation network task. ACL 2013, 153 (2013)

25 BioCreAtivE: Critical Assessment of Information Extraction in Biology. http://www.biocreative.org

26 NaCTeM: The National Centre for Text Mining. http://www.nactem.ac.uk/

27 CALBC: Collaborative Annotation of a Large Biomedical Corpus. http://goo.gl/5VUP2u

28 Hakenberg: Icahn School of Medicine at Mount Sinai Hospital. http://goo.gl/EhkmZm

29 Berlin: Institut für Informatik at Humboldt-Universität zu Berlin. http://corpora.informatik.hu-berlin.de/

30 Kim, J.-D., Nguyen, N., Wang, Y., Tsujii, J., Takagi, T., Yonezawa, A.: The genia event and protein coreference tasks of the bionlp shared task 2011. BMC bioinformatics 13(Suppl 11), 1 (2012)

31 Ohta, T., Pyysalo, S., Tsujii, J.: Overview of the epigenetics and post-translational modifications (epi) task of bionlp shared task 2011. In: Proceedings of the BioNLP Shared Task 2011 Workshop, pp. 16–25 (2011). Association for Computational Linguistics

32 Pyysalo, S., Ohta, T., Tsujii, J.: Overview of the entity relations (rel) supporting task of bionlp shared task 2011. In: Proceedings of the BioNLP Shared Task 2011 Workshop, pp. 83–88 (2011). Association for Computational Linguistics

33 Pyysalo, S., Ohta, T., Ananiadou, S.: Overview of the cancer genetics (cg) task of bionlp shared task 2013. In: Proceedings of the BioNLP Shared Task 2013 Workshop, pp. 58–66 (2013). Association for Computational Linguistics

34 Ohta, T., Pyysalo, S., Rak, R., Rowley, A., Chun, H.-W., Jung, S.-J., Jeong, C.-h., Choi, S.-p., Ananiadou, S.: Overview of the pathway curation (pc) task of bionlp shared task 2013. ACL 2013 (2013). Association for Computational Linguistics

35 Kim, J.-j., Han, X., Lee, V.: Gro task: Populating the gene regulation ontology with events and relations. ACL 2013, 50 (2013)

36 Björne, J., Salakoski, T.: Tees 2.1: Automated annotation scheme learning in the bionlp 2013 shared task. ACL 2013, 16 (2013). Association for Computational Linguistics

37 Björne, J., Ginter, F., Salakoski, T.: University of turku in the bionlp’11 shared task. BMC bioinformatics 13(Suppl 11), 4 (2012)

38 Tsochantaridis, I., Joachims, T., Hofmann, T., Altun, Y., Singer, Y.: Large margin methods for structured and interdependent output variables. Journal of Machine Learning Research 6(9) (2005)

39 McClosky, D., Charniak, E.: Self-training for biomedical parsing. In: Proceedings of the 46th Annual Meeting of the ACL on Human Language Technologies: Short Papers, pp. 101–104 (2008). Association for Computational Linguistics

40 De Marneffe, M.-C., Manning, C.D.: The stanford typed dependencies representation. In: Coling 2008: Proceedings of the Workshop on Cross-Framework and Cross-Domain Parser Evaluation, pp. 1–8 (2008). Association for Computational Linguistics

41 Björne, J., Airola, A., Pahikkala, T., Salakoski, T.: Drug-drug interaction extraction from biomedical texts with svm and rls classifiers. Proceedings of DDIExtraction-2011 challenge task, 35–42 (2011)

42 Björne, J., Kaewphan, S., Salakoski, T.: Uturku: Drug named entity recognition and drug-drug interaction extraction using svm classification and domain knowledge. In: Proceedings of the Seventh International Workshop on Semantic Evaluation, pp. 651–659 (2013). Association for Computational Linguistics

43 Björne, J., Ginter, F., Pyysalo, S., Tsujii, J., Salakoski, T.: Complex event extraction at pubmed scale. Bioinformatics 26(12), 382–390

44 Van Landeghem, S., Björne, J., Wei, C.-H., Hakala, K., Pyysalo, S., Ananiadou, S., Kao, H.-Y., Lu, Z., Salakoski, T., Van de Peer, Y., et al.: Large-scale event extraction from literature with multi-level gene normalization. PloS one 8(4), 55814 (2013)

45 Neves, M., Damaschun, A., Mah, N., Lekschas, F., Seltmann, S., Stachelscheid, H., Fontaine, J.-F., Kurtz, A., Leser, U.: Preliminary evaluation of the cellfinder literature curation pipeline for gene expression in kidney cells and anatomical parts. Database: the journal of biological databases and curation (2013)

46 Gerner, M., Sarafraz, F., Bergman, C.M., Nenadic, G.: Biocontext: an integrated text mining system for large-scale extraction and contextualization of biomolecular events. Bioinformatics 28(16), 2154–2161 (2012)

47 Szostak, J., Ansari, S., Madan, S., Fluck, J., Talikka, M., Iskandar, A., De Leon, H., Hofmann-Apitius, M., Peitsch, M.C., Hoeng, J.: Construction of biological networks from unstructured information based on a semi-automated curation workflow. Database 2015, 057 (2015)

48 Miwa, M., Ananiadou, S.: Nactem eventmine for bionlp 2013 cg and pc tasks. In: Proceedings of BioNLP Shared Task 2013 Workshop, pp. 94–98 (2013)

49 Miwa, M., Ananiadou, S.: Adaptable, high recall, event extraction system with minimal configuration. BMC bioinformatics 16(Suppl 10), 7 (2015)

50 Fan, R.-E., Chang, K.-W., Hsieh, C.-J., Wang, X.-R., Lin, C.-J.: Liblinear: A library for large linear classification. The Journal of Machine Learning Research 9, 1871–1874 (2008)

51 Sagae, K., Tsujii, J.: Dependency parsing and domain adaptation with lr models and parser ensembles. In: Proceedings of the CoNLL Shared Task Session of EMNLP-CoNLL, pp. 1044–1050 (2007). Association for Computational Linguistics

52 Miyao, Y., Sagae, K., Sætre, R., Matsuzaki, T., Tsujii, J.: Evaluating contributions of natural language parsers to protein-protein interaction extraction. Bioinformatics 25(3), 394–400 (2009)

53 Miwa, M., Thompson, P., Ananiadou, S.: Boosting automatic event extraction from the literature using domain adaptation and coreference resolution. Bioinformatics 28(13), 1759–1765 (2012)

54 Miwa, M., Thompson, P., McNaught, J., Kell, D.B., Ananiadou, S.: Extracting semantically enriched events from biomedical literature. BMC bioinformatics 13(1), 108 (2012)

55 McClosky, D., Surdeanu, M., Manning, C.D.: Event extraction as dependency parsing. In: Proceedings of the 49th Annual Meeting of the ACL: Human Language Technologies - Volume 1, pp. 1626–1635 (2011). Association for Computational Linguistics

56 McDonald, R., Lerman, K., Pereira, F.: Multilingual dependency analysis with a two-stage discriminative parser. In: Proceedings of the Tenth Conference on CoNLL, pp. 216–220 (2006). Association for Computational Linguistics

57 Riedel, S., McCallum, A.: Fast and robust joint models for biomedical event extraction. In: Proceedings of the Conference on EMNLP, pp. 1–12 (2011). Association for Computational Linguistics

58 Zhang, Z., Iria, J., Ciravegna, F.: Improving domain-specific entity recognition with automatic term recognition and feature extraction. In: Proceedings of the Seventh Conference on International Language Resources and Evaluation (2010). European Languages Resources Association

59 Bui, Q.-C., Van Mulligen, E.M., Campos, D., Kors, J.A.: A fast rule-based approach for biomedical event extraction. ACL 2013, 104 (2013)

60 Bui, Q.-C., Sloot, P.M., van Mulligen, E.M., Kors, J.A.: A novel feature-based approach to extract drug-drug interactions from biomedical text. Bioinformatics, 557 (2014)

61 Cohen, K.B., Verspoor, K., Johnson, H.L., Roeder, C., Ogren, P.V., Baumgartner Jr, W.A., White, E., Tipney, H., Hunter, L.: High-precision biological event extraction with a concept recognizer. In: Proceedings of the Workshop on Current Trends in BioNLP: Shared Task, pp. 50–58 (2009). Association for Computational Linguistics

62 Kilicoglu, H., Bergler, S.: Adapting a general semantic interpretation approach to biological event extraction. In: Proceedings of the BioNLP Shared Task 2011 Workshop, pp. 173–182 (2011). Association for Computational Linguistics

63 Kilicoglu, H., Bergler, S.: Syntactic dependency based heuristics for biological event extraction. In: Proceedings of the Workshop on Current Trends in BioNLP: Shared Task, pp. 119–127 (2009). Association for Computational Linguistics

64 Kang, N., Singh, B., Afzal, Z., van Mulligen, E.M., Kors, J.A.: Using rule-based natural language processing to improve disease normalization in biomedical text. Journal of the American Medical Informatics Association, 2012 (2012)

65 Allen, J., de Beaumont, W., Galescu, L., Teng, C.M.: Complex event extraction using drum. In: Proceedings of BioNLP 15, pp. 1–11 (2015). Association for Computational Linguistics

66 Liu, H., Verspoor, K., Comeau, D.C., MacKinlay, A., Wilbur, W.J.: Generalizing an approximate subgraph matching-based system to extract events in molecular biology and cancer genetics. ACL 2013, 76 (2013)

67 Airola, A., Pyysalo, S., Björne, J., Pahikkala, T., Ginter, F., Salakoski, T.: All-paths graph kernel for protein-protein interaction extraction with evaluation of cross-corpus learning. BMC bioinformatics 9(Suppl 11), 2 (2008)

68 Van Landeghem, S., Björne, J., Abeel, T., De Baets, B., Salakoski, T., Van de Peer, Y.: Semantically linking molecular entities in literature through entity relationships. BMC bioinformatics 13(Suppl 11), 6 (2012)

69 Domingos, P., Lowd, D.: Markov logic: An interface layer for artificial intelligence. Synthesis Lectures on Artificial Intelligence and Machine Learning 3(1), 1–155 (2009)

70 Richardson, M., Domingos, P.: Markov logic networks. Machine learning 62(1–2), 107–136 (2006)

71 Riedel, S., Chun, H.-W., Takagi, T., Tsujii, J.: A markov logic approach to bio-molecular event extraction. In: Proceedings of the Workshop on Current Trends in BioNLP: Shared Task, pp. 41–49 (2009). Association for Computational Linguistics

72 Taskar, B., Wong, M.-F., Abbeel, P., Koller, D.: Link prediction in relational data. In: Advances in Neural Information Processing Systems, p. 8 (2003)

73 Poon, H., Vanderwende, L.: Joint inference for knowledge extraction from biomedical literature. In: Human Language Technologies: The 2010 Annual Conference of the North American Chapter of the ACL, pp. 813–821 (2010). Association for Computational Linguistics

74 Kok, S., Singla, P., Richardson, M., Domingos, P., Sumner, M., Poon, H., Lowd, D.: The alchemy system for statistical relational ai. University of Washington (2005)

75 Venugopal, D., Chen, C., Gogate, V., Ng, V.: Relieving the computational bottleneck: Joint inference for event extraction with high-dimensional features. In: Proceedings of the Conference on EMNLP, pp. 831–843 (2014). Association for Computational Linguistics

76 Riedel, S.: Improving the accuracy and efficiency of map inference for markov logic. In: Proceedings of the 24th Conference on Uncertainty in Artificial Intelligence, UAI 2008, pp. 468–475 (2008)

77 Riedel, S., Sætre, R., Chun, H.-W., Takagi, T., Tsujii, J.: Bio-molecular event extraction with markov logic. Computational Intelligence 27(4), 558–582 (2011)

78 Rush, A.M., Collins, M.: A tutorial on dual decomposition and lagrangian relaxation for inference in natural language processing. Journal of Artificial Intelligence Research 45, 305–362 (2012)

79 Riedel, S., McCallum, A.: Robust biomedical event extraction with dual decomposition and minimal domain adaptation. In: Proceedings of the BioNLP Shared Task 2011 Workshop, pp. 46–50 (2011). Association for Computational Linguistics

80 Riedel, S., McClosky, D., Surdeanu, M., McCallum, A., Manning, C.D.: Model combination for event extraction in bionlp 2011. In: Proceedings of the BioNLP Shared Task 2011 Workshop, pp. 51–55 (2011). Association for Computational Linguistics

81 Liu, X., Bordes, A., Grandvalet, Y.: Extracting biomedical events from pairs of text entities. BMC bioinformatics 16(Suppl 10), 8 (2015)

82 Vlachos, A., Craven, M.: Search-based structured prediction applied to biomedical event extraction. In: Proceedings of the Fifteenth Conference on CoNLL, pp. 49–57 (2011). Association for Computational Linguistics

83 Liu, X., Bordes, A., Grandvalet, Y., et al.: Biomedical event extraction by multi-class classification of pairs of text entities. In: Proceedings of the BioNLP Shared Task 2013 Workshop, pp. 45–49 (2013)

84 Kordjamshidi, P., Roth, D., Moens, M.-F.: Structured learning for spatial information extraction from biomedical text: bacteria biotopes. BMC bioinformatics 16(1), 129 (2015)

85 Collins, M.: Discriminative training methods for hidden markov models: Theory and experiments with perceptron algorithms. In: Proceedings of the ACL-02 Conference on Empirical Methods in Natural Language processing-Volume 10, pp. 1–8 (2002). Association for Computational Linguistics

86 OntoBiotope: OntoBiotope habitat ontology. http://goo.gl/JdBR2Z

87 NCBI: NCBI bacterium taxonomy database. http://www.ncbi.nlm.nih.gov/taxonomy

88 Kordjamshidi, P., Moens, M.-F.: Designing constructive machine learning models based on generalized linear learning techniques. In: Proceedings of the NIPS Workshop on Constructive Machine Learning, pp. 1–5 (2013)

89 EVEX: A PubMed-Scale Resource for Homology-Based Generalization of Text Mining Predictions. http://evexdb.org/

90 FACTA+: Finding Associated Concepts with Text Analysis. http://www.nactem.ac.uk/facta/

91 McClosky, D., Charniak, E., Johnson, M.: Automatic domain adaptation for parsing. In: Human Language Technologies: The 2010 Annual Conference of the North American Chapter of the ACL, pp. 28–36 (2010). Association for Computational Linguistics

92 Buyko, E., Hahn, U.: Evaluating the impact of alternative dependency graph encodings on solving event extraction tasks. In: Proceedings of the 2010 Conference on Empirical Methods in Natural Language Processing, pp. 982–992 (2010). Association for Computational Linguistics

93 Garten, Y., Coulet, A., Altman, R.B.: Recent progress in automatically extracting information from the pharmacogenomic literature. Pharmacogenomics 11(10), 1467–1489 (2010)

94 Hirst, G.: Ontology and the lexicon. In: Staab, S., Studer, R. (eds.) Handbook on Ontologies. International Handbooks on Information Systems, pp. 269–292. Springer, ??? (2009)

95 Jimeno-Yepes, A., Jimenez-Ruiz, E., Berlanga-Llavori, R., Rebholz-Schuhmann, D.: Reuse of terminological resources for efficient ontological engineering in life sciences. BMC bioinformatics 10(Suppl 10), 4 (2009)

96 Buyko, E., Beisswanger, E., Hahn, U.: Testing different ace-style feature sets for the extraction of gene regulation relations from medline abstracts. In: Proceedings of the Third International Symposium on Semantic Mining in Biomedicine (SMBM), pp. 21–28 (2008). TUCS

97 Johnson, H.L., Baumgartner, W.A., Krallinger, M., Cohen, K.B., Hunter, L.: Corpus refactoring: a feasibility study. Journal of Biomedical Discovery and Collaboration 2(1), 4 (2007)

98 Lavergne, T., Grouin, C., Zweigenbaum, P.: The contribution of co-reference resolution to supervised relation detection between bacteria and biotopes entities. BMC bioinformatics 16(Suppl 10), 6 (2015)

99 Zerva, C., Ananiadou, S.: Event extraction in pieces: Tackling the partial event identification problem on unseen corpora, 31–41 (2015)

100 LeCun, Y., Bengio, Y., Hinton, G.: Deep learning. Nature 521(7553), 436–444 (2015)

101 Li, C., Song, R., Liakata, M., Vlachos, A., Seneff, S., Zhang, X.: Using word embedding for bio-event extraction. In: Proceedings of BioNLP 15, pp. 121–126 (2015). Association for Computational Linguistics

102 Spranger, M., Palaniappan, S.K., Rennes, F., Ghosh, S.: Extracting biological pathway models from nlp event representations. ACL-IJCNLP 2015, 42 (2015)

